# Theta gates and routes information in the frontal cortex

**DOI:** 10.64898/2026.06.03.729810

**Authors:** Matthew B. Broschard, Scott L. Brincat, Roman F. Loonis, Earl K. Miller

## Abstract

Theta (4–10 Hz) oscillations seem well-suited for coordinating neural activity. Many studies have focused on theta’s role in long-range coordination across brain regions (e.g., connectivity between the prefrontal cortex and the hippocampus). It remains unclear how theta coordinates neural activity more locally within prefrontal subareas. We examined neural activity in three frontal areas (i.e., dorsolateral prefrontal cortex (dlPFC), ventrolateral prefrontal cortex (vlPFC), and frontal eye fields (FEF)) as non-human primates categorized dot patterns. We found that theta flexibly coordinated spiking activity and higher-frequency oscillations within and between frontal areas. First, theta phase in all areas was coupled to spiking information in the FEF and seemed to gate information that was behaviorally relevant. Second, theta influences were routed in opposite directions depending on feedback. Theta flowed in a posterior direction to the FEF during choices and after correct outcomes. Theta influences reversed directionality and flowed in an anterior direction after incorrect outcomes. Third, theta organized nested cross-frequency, phase-amplitude interactions. Theta was coupled to beta (15-30Hz) oscillations, both within and between areas. Beta, in turn, was coupled to gamma (40-90Hz) oscillations, but mainly locally. Together, our results position theta as a critical mechanism that flexibly and dynamically coordinates neural activity within and across the frontal cortex.

**HIGHLIGHTS:**

- Neural activity was recorded in three frontal areas as non-human primates categorized dot patterns.
- Theta (4-10Hz) oscillations gated relevant information in spiking.
- Theta influences were routed in opposite directions depending on feedback.
- Theta organized nested cross-frequency interactions with beta (15-30Hz) and gamma (40-90Hz) oscillations.

**Graphical Abstract:** 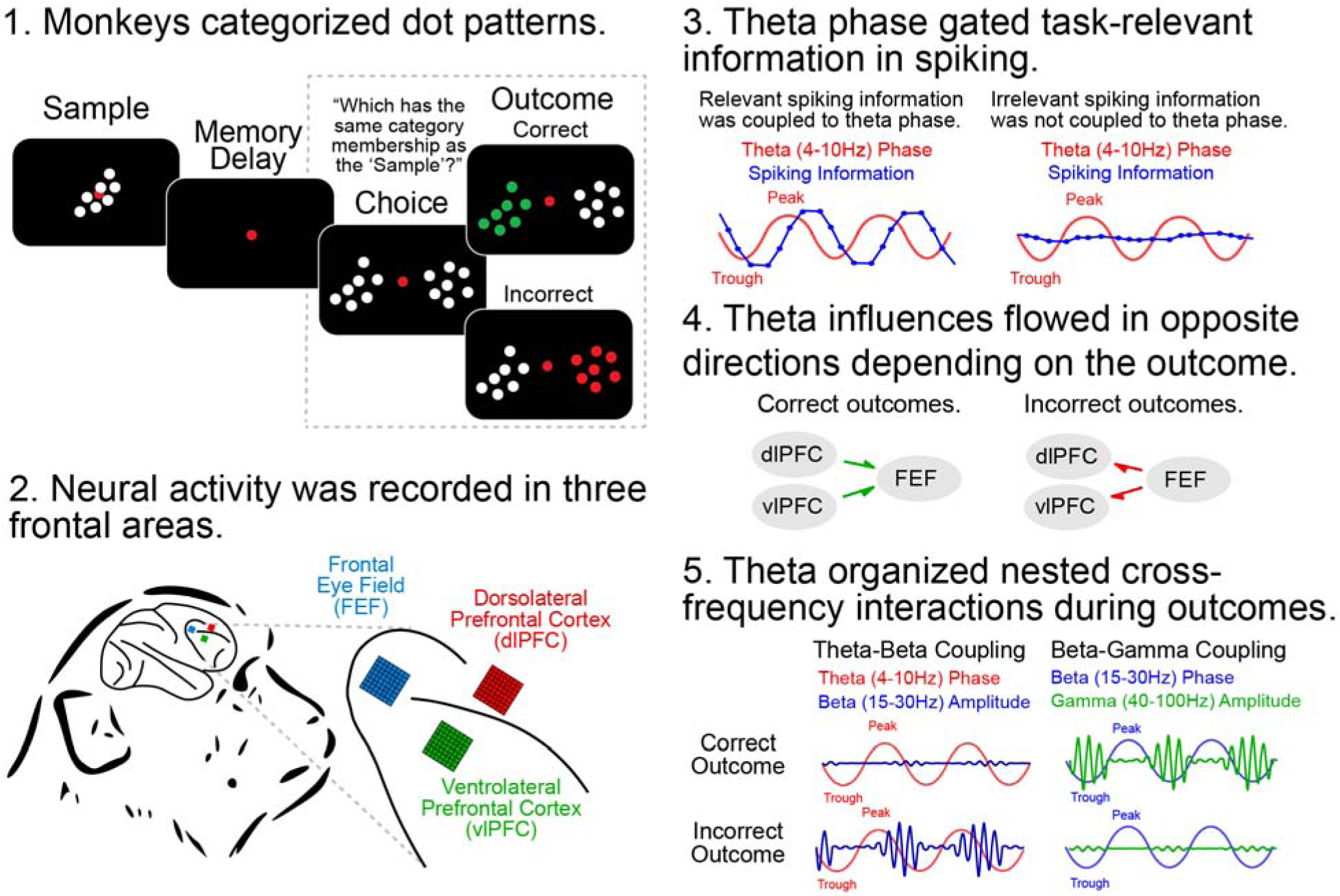

## INTRODUCTION

Cognition is not static. The brain must dynamically shift how information is coordinated and updated within and between regions. This flexibility seems to be achieved, in part, by neural oscillations (Buzsáki & Draguhn, 2004; Engel, Fries, & Singer, 2001; Fries, 2015). Oscillations organize neural activity within local circuits (Lei et al., 2026; Cardin et al., 2009; Lundqvist et al., 2016) and across larger distances (Gregoriou, Gotts, Zhou, & Desimone, 2009; Buschman & Miller, 2007; von Stein & Sarnthein, 2000).

Theta (4–10 Hz) oscillations, in particular, seem to be well suited for coordinating neural activity. Theta oscillations synchronize regions across long-range distances (Siapas et al., 2005; Jones & Wilson, 2005; Fujisawa & Buzsáki 2011; Liebe et al., 2012). Theta organizes into large-scale traveling waves across the cortex (Zhang, Watrous, Patel, & Jacobs, 2018; Patel et al., 2012). And theta routes information selectively through distinct pathways (Colgin et al., 2009; Colgin, 2013; Hasselmo et al., 2002; Helfrich et al., 2018; Fiebelkorn & Kastner, 2019). Within the prefrontal cortex, theta oscillations tend to increase when cognitive control is needed (Womelsdorf, Vinck, Leung, & Everling, 2010; Han et al., 2026). Theta is observed during surprising events, such as negative feedback (Cavanagh, Frank, Klein, & Allen, 2010; Trujillo & Allen, 2007; van de Vijver et al., 2011; Andreou et al., 2017; Brincat & Miller, 2015; Loonis et al., 2017) and stimulus conflict (Muralidharan et al., 2023; Choi et al., 2024). Furthermore, theta synchrony peaks around “decision-points” (Cavanagh & Frank, 2014) to integrate information from multiple sources (e.g., task rules, stimulus-reward associations, and reward expectations; Womelsdorf, Johnston, Vinck, & Everling, 2010; Benchenane et al., 2010; van Wingerden et al., 2010).

The majority of these studies have focused on a single role for theta in isolation. Here, we address whether theta can dynamically shift its role according to changing task demands, thus supporting the flexibility associated with cognitive function. Additionally, the coordinating role of theta in the prefrontal cortex has largely focused on long-range connectivity (e.g., with hippocampus or visual cortex). It is generally underexplored how theta coordinates processing within the frontal cortex. Thus, we examined neural activity from multiple frontal areas, simultaneously.

We examined the neural dynamics as non-human primates (NHPs) categorized dot patterns. Neural activity was recorded in three frontal areas: the dorsolateral prefrontal cortex (dlPFC), the ventrolateral prefrontal cortex (vlPFC), and the frontal eye fields (FEF). We found that the FEF emerged as a critical hub during decision-making and feedback processing. Theta phase, in all areas, was selectively coupled to FEF spiking information. The type of information coupled to theta shifted across the trial and depended on what was immediately relevant. This was accompanied with a change in the directionality of theta causality. Theta influences flowed in the posterior direction toward the FEF during choices and after correct outcomes. By contrast, theta influences flowed in the anterior direction away from the FEF after incorrect outcomes. Phase-amplitude coupling revealed that theta phase organized higher-frequency oscillations. Theta was coupled to beta oscillations, within and across areas. Beta, in turn, was coupled to gamma oscillation, more locally.

## RESULTS

Non-human primates (NHPs) learned to categorize dot patterns (Fig. 1A; Loonis et al., 2017; Posner & Keele, 1968). Two new dot patterns were generated before each session and functioned as the category prototypes (see Methods). The NHPs never saw these prototypes. Instead, they were shown category exemplars generated by randomly distorting the dot positions of the prototypes. During each trial, a category exemplar appeared at the center of the screen (Fig. 1B; sample epoch). After a memory delay (delay epoch), two new exemplars were shown on the left and right sides of the screen (choice epoch). One of the choice exemplars came from the same category as the sample exemplar presented earlier (i.e., its category “match”). The other exemplar came from the other category (category “non-match”). The NHPs were trained to fixate on the “match” exemplar to receive a reward (correct choice). Visual feedback was given simultaneously with the reward (outcome). After correct choices, the color of the chosen exemplar turned from white to green. After incorrect choices, the color of the chosen exemplar turned red, and a “timeout” was given before the start of the next trial.

**Figure 1.**
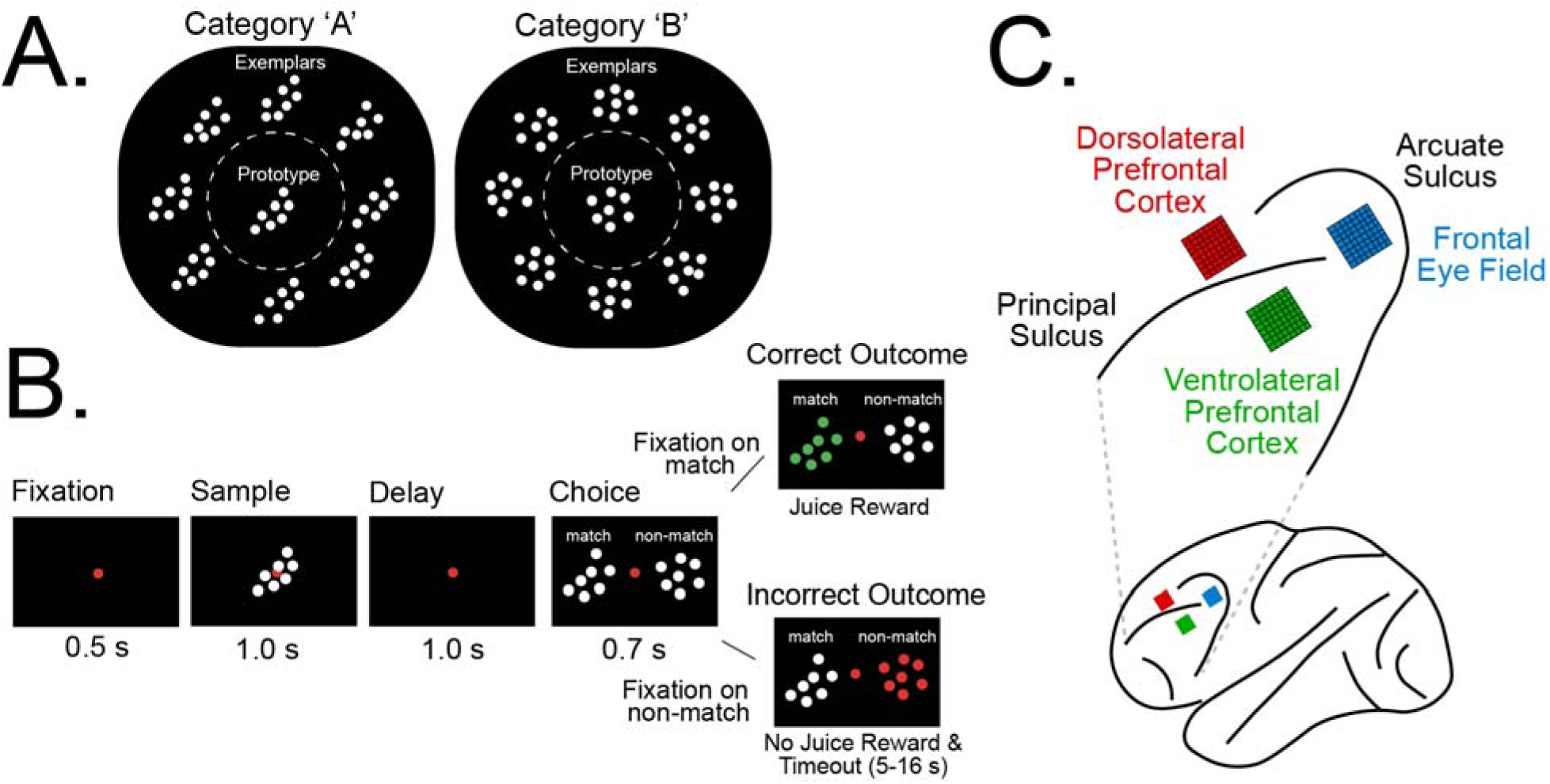
NHPs learned to categorize dot patterns. **A,** Two “prototype” dot patterns were generated before each session. Category exemplars were generated by distorting the dot positions of these prototypes. **B,** Trial procedure. After fixation, a category exemplar was presented at the center of the screen (sample). After a memory delay (delay), two new exemplars appeared on the left and right sides of the screen (choice). The NHPs were trained to fixate on the exemplar that came from the same category as the sample (i.e., category “match”). After correct choices, the chosen exemplar turned from white to green, and a juice reward was delivered (correct outcome). After incorrect choices (i.e., the NHPs fixated on the other, category “non-match”), the chosen exemplar turned red, and a timeout was given without juice (i.e., incorrect outcome). **C,** The approximate location of the electrode arrays centered in the dorsolateral prefrontal cortex (dlPFC), ventrolateral prefrontal cortex (vlPFC), and the frontal eye fields (FEF).

Local field potentials (LFPs) and spiking activity were recorded from three electrode arrays positioned in the frontal cortex (Fig 1C). Two arrays were centered in more anterior locations in the dorsolateral prefrontal cortex (dlPFC) and the ventrolateral prefrontal cortex (vlPFC). The third array was centered in a more posterior location in the frontal eye fields (FEF). The current analysis only included sessions in which the NHPs successfully learned the categories (see Methods; 66 out of 76 total sessions; average number of trials: 777.48 ± 219.21 SD; average accuracy: 78.10% ± 5.73 SD). For this analysis, we examined trials after the categories were learned (see Methods) and focused on neural activity during the choice and outcome trial epochs.

### Spiking information and LFP power reflected choices and outcomes

Spiking activity in all areas carried information about task variables. We trained Support Vector Machine (SVM) classifiers to decode information about three task variables: the sample’s category membership (category information; category ‘A’ vs. category ‘B’), the direction of the correct choice exemplar (choice information; left vs. right), and the valence of the outcome (outcome information; correct vs. incorrect). For each variable, classifier accuracy increased above chance level (50%) after information about that variable became available within the trial (Fig. 2A & Supplemental Fig. 1). Category information increased after the sample exemplar was shown (Supplemental Fig. 1), choice information increased after the choice exemplars were shown (Fig. 2A), and outcome information increased after feedback was given (Fig. 2A). Information about each variable remained above chance throughout the remainder of the trial. Thus, spiking activity carried information about multiple task variables simultaneously during the choice and outcome epochs. These patterns were observed in all three areas and were consistent for each NHP subject (Supplemental Fig. 1). However, spiking information tended to be stronger in the vlPFC and the FEF than the dlPFC, for all variables (Fig. 2A).

**Figure 2.**
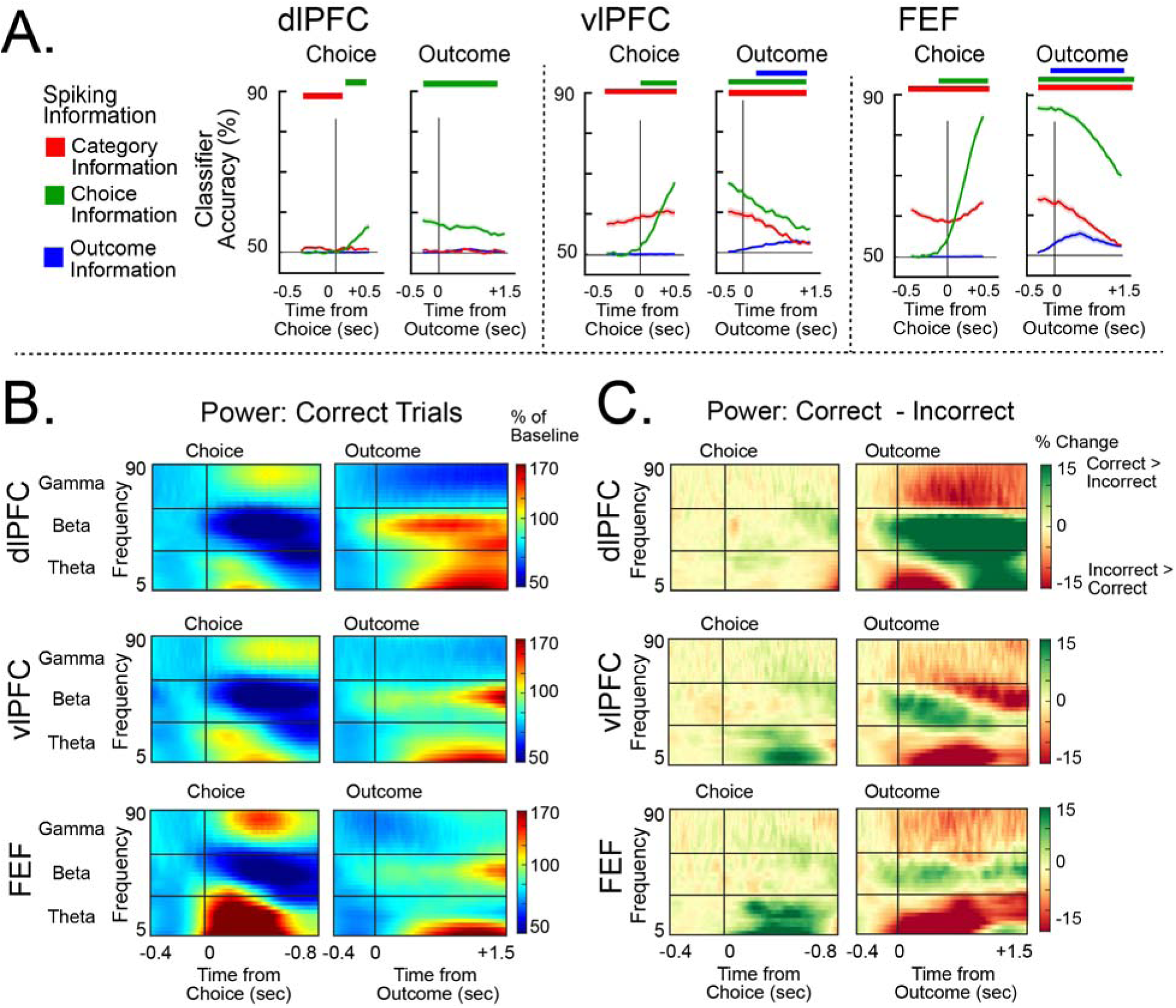
Spiking information and LFP power reflected choices and outcomes. **A,** Support Vector Machine (SVM) classifiers were trained with spiking activity from each area to decode the following task information: category (category ‘A’ vs. category ‘B’; red), choice (‘left’ vs. ‘right’ behavioral response; green), and trial outcome (‘correct’ vs. ‘incorrect’; blue). Colored bars above each plot indicate timepoints in which classifier accuracy was significantly above chance level (50%; permutation test, *p* < .05). All error bars indicate *S.E.M.* **B,** Heatmaps of average LFP power around choices (left) and outcomes (right) during correct trials. Horizontal lines indicate the theta, beta, and gamma frequency bands. **C,** Heatmaps of difference in average LFP power (correct trials – incorrect trials) around choices (left) and outcomes (right). The color indicates whether power was stronger during correct trials (green) or incorrect trials (red). All error bars indicate *S.E.M*.

LFP power also reflected choices and outcomes. We examined LFP power, averaged in the theta (4–10 Hz), beta (15–30 Hz), and gamma (40–90 Hz) frequency bands (Figs. 2B&C & Supplemental Fig. 2). During the sample and delay epochs, LFP power showed relatively modest changes and did not differ between correct and incorrect trials (Supplemental Fig. 2). By contrast, LFP power was more strongly modulated during the choice and outcome epochs, and power differed between correct and incorrect trials. During choices, theta and gamma power increased, and beta power decreased, in all areas (Fig. 2B, left). Theta power was stronger during correct trials than incorrect trials in the vlPFC and the FEF (Fig. 2C & Supplemental Fig. 2A; vlPFC: Cohen’s *d* = 0.53; FEF: *d =* 0.50) but not the dlPFC (Fig. 2C & Supplemental Fig. 2A; Cohen’s *d* = 0.01), suggesting that theta power in the vlPFC/FEF was important for successful behavior. During the outcome epoch, theta and beta power increased, and gamma power decreased (Fig. 2B, right). For all areas, theta and gamma power were stronger after incorrect outcomes than correct outcomes (Fig. 2C & Supplemental Fig. 2; Theta. dlPFC: Cohen’s *d* = -0.27; vlPFC: *d* = -0.26; FEF: *d* = -0.45. Gamma: dlPFC: *d* = -2.61; vlPFC: *d* = - 0.56; FEF: *d* = -1.54). By contrast, beta power was stronger after correct outcomes than incorrect outcomes (Fig. 2C & Supplemental Fig. 2; dlPFC: *d* = 1.62; vlPFC: *d =* 0.48; FEF: *d* = 0.74).

### Spiking information varied with theta phase

Above, we found that spiking activity and LFP power reflected choices and their outcomes. Next, we tested whether information carried by spiking varied with the phase of the ongoing LFP. For each LFP frequency, we divided LFP phase (averaged across all electrodes in an array) into 45-degree bins. Then, SVM classifiers were trained with spiking activity from each phase bin, and phase-locking values quantified whether classifier accuracies varied across phase bins. We examined within-area comparisons, where the spiking activity and the LFP phase were computed from the same array. We also examined between-area comparisons, where spiking activity and LFP phase were computed from different arrays.

Category, choice, and outcome information in spiking were coupled to LFP theta phase, but at different times during the trial. To preview the results, these effects only occurred with FEF spiking activity during the choice and outcome epochs, but with both local (within-area) and distant (between-area) LFP theta phase. During the choice epoch, choice information in FEF spiking was coupled to local FEF theta phase (Fig. 3A, left; Cohen’s *d* = 0.98) and distant theta phase in the dlPFC and vlPFC (Fig. 3B, left; dlPFC theta: *d* = 0.87; vlPFC theta: *d* = 0.90). FEF choice information was strongest at the falling phases of theta and weakest at the rising phases of theta, both for local FEF theta phase (Fig. 3C, left; preferred phase = 80.67 degrees ± 9.5 *SEM*) and distant dlPFC/vlPFC theta phase (Fig. 3D, left; preferred dlPFC phase = 104.40 degrees ± 10.3 *SEM*; preferred vlPFC phase = 78.59 degrees ± 11.4 *SEM*). These patterns were consistent for both NHPs (Supplemental Figs. 3D&E). Also during the choice epoch, category information in FEF spiking was coupled to local FEF theta phase (Fig. 3A, left; Cohen’s *d* = 0.33). However, this was only observed for one of the NHPs (Supplemental Fig. 3D). During the outcome epoch, FEF choice information and category information were no longer coupled to theta phase (FEF choice information: Cohen’s *d* = 0.07; FEF category information: *d* = -0.02), even though average spiking activity still carried information about them (Fig. 2A). Instead, outcome information in FEF spiking was coupled to FEF theta phase (Fig. 3A, right; Cohen’s *d* = 0.59) and distant dlPFC/vlPFC theta phase (Fig. 3B, right; dlPFC theta: *d* = 0.72; vlPFC theta: *d* = 0.58). Like during the choice epoch, FEF outcome information was strongest during the falling phases of theta and weakest during the rising phases of theta, for local FEF theta phase (Fig. 3C, right; preferred phase = 99.58 degrees ± 10.7 *SEM*) and distant dlPFC/vlPFC theta phase (Fig. 3D, right; preferred dlPFC phase = 103.80 degrees ± 11.2 *SEM*; preferred vlPFC theta phase = 130.52 degrees ± 10.9 *SEM*). These patterns were consistent for both NHPs (Supplemental Figs. 3D&E). All other comparisons were not significant (Supplemental Figs. 3A-C).

**Figure 3.**
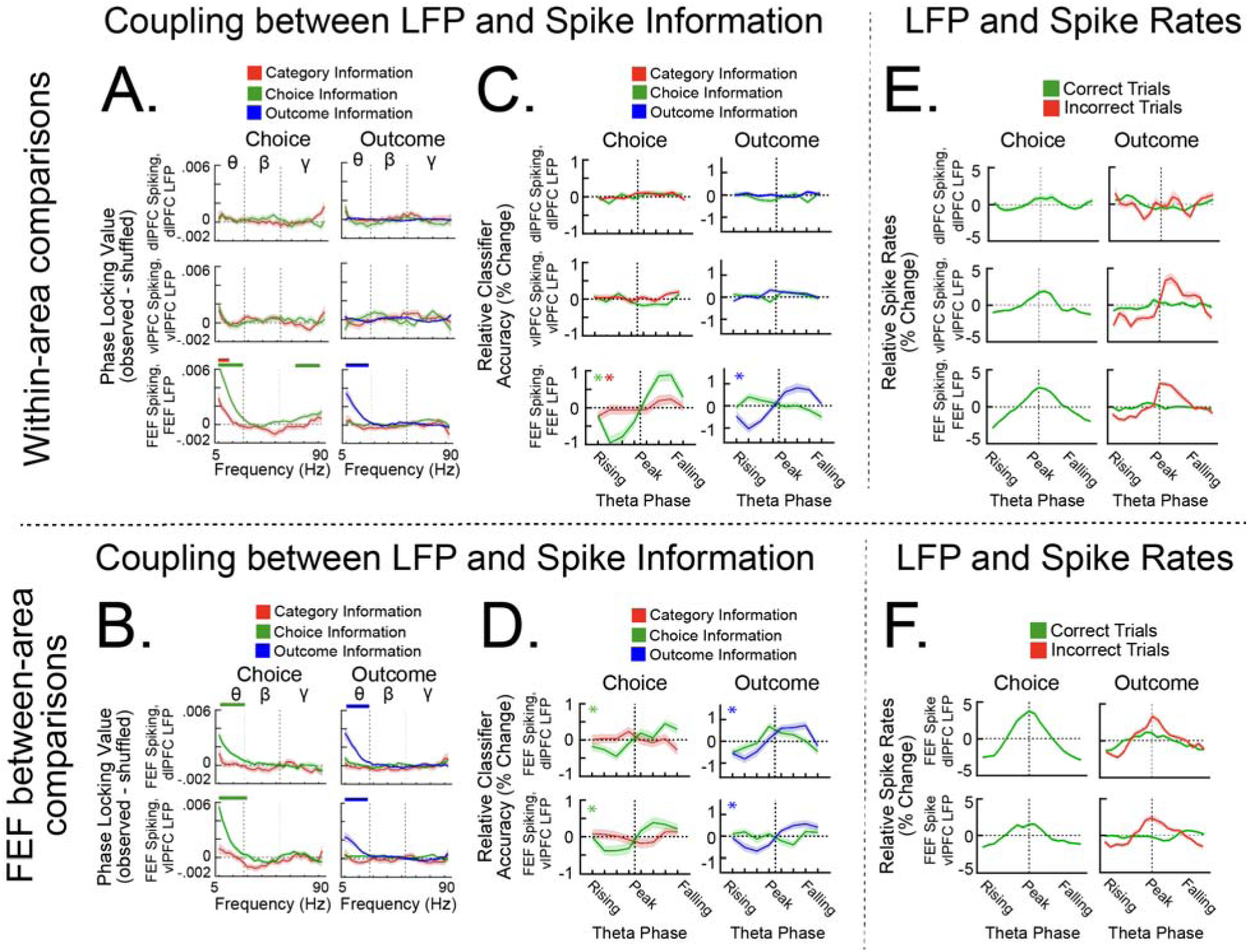
FEF spiking information was coupled to theta phase during choices and outcomes **A-B,** LFP phase was divided into bins, and SVM classifiers were trained to decode task information using spiking activity from each phase bin. Phase-locking values quantified whether spiking information varied with LFP phase. Colored bars above each plot indicate LFP frequencies that were significantly coupled to spiking information (permutation test *p* < .05). Supplemental Fig. 3A-C shows the results during the sample and delay epochs, as well as for all within-area and between-area comparisons. **A,** Coupling between spiking information and LFP phase for all within-area comparisons, where spiking activity and LFP phase were computed from the same array. **B,** Coupling between FEF spiking activity and distant LFP phase in the dlPFC and the vlPFC. **C-D,** Spiking information plotted across theta phase bins for the results in (**A-B**). * in the top left quadrant of each plot (and its color) indicates the type of information that was significantly coupled to theta phase during that trial epoch. **C,** Within-area spiking information across theta phase bins (same comparisons as panel **A**). **D,** Between-area FEF spiking information across theta phase bins (same comparisons as panel **B**). **E-F,** Overall spike rates, (rather than spiking information), across theta phase bins. **E,** Within-area spike rates across theta phase bins (same comparisons as panels **A** and **C**). **F,** Between-area FEF spike rates across theta phase bins (same comparisons as panels **B** and **D**). All error bars indicate *S.E.M.* θ, β, γ refer to theta, beta, and gamma frequency bands, respectively.

The optimal phase of decoding spiking information was offset from the preferred phase of overall spike-field coupling. Above, we found that FEF spiking information was strongest during the falling phases of theta. We examined whether this was also true for overall spikes rates by binning spiking activity across theta phases. We found that spike rates were highest during the peak phases of theta. This was consistent for within-area comparisons (Fig. 3E; mean preferred phase = 5.1 degrees ± 13.1 *SEM*), between-area comparisons (Fig. 3F; Supplemental Fig. 4; mean preferred phase = 2.5 degrees ± 9.2 *SEM*), and across all trial epochs (Supplemental Fig. 4). Thus, overall spike rates tended to peak earlier in phase than spiking information.

### Theta influences flowed in opposite directions during choices and outcomes

The results above indicate that theta phase, in all areas, was coupled to FEF spiking information. The finding that theta coupled specifically to FEF spiking motivated us to look at how theta flowed across the subareas. Thus, we computed Granger causality to examine the directionality of theta influences. For this analysis, and all following analyses, we focused on the choice and outcome epochs, when spiking information was coupled to theta phase (see above).

Theta influences flowed in opposite directions during choices and outcomes. During the choice epoch, theta influences flowed in the posterior direction from the dlPFC and the vlPFC to the FEF (Fig. 4A; dlPFC → FEF directionality: Cohen’s *d* = 0.94; vlPFC → FEF directionality: *d* = 0.41). This was consistent for both NHPs (Supplemental Fig. 5A). Beta and gamma causality did not show directionality at this time (Supplemental Figs. 5B&C). During the outcome epoch, the directionality of theta causality depended on whether the choice was correct or incorrect. After correct choices, theta influences continued to flow posteriorly from the dlPFC and the vlPFC to the FEF (Fig. 4B; dlPFC → FEF directionality: Cohen’s *d* = 0.49; vlPFC → FEF directionality: *d* = 0.32). However, after incorrect choices, theta directionality reversed and was stronger from the FEF to the dlPFC and the vlPFC (Fig. 4B; FEF → dlPFC directionality: *d* = - 0.56; FEF → vlPFC directionality: *d* = -0.67; Fig. 4C summarizes these effects). This was consistent for both NHPs (Supplemental Fig. 5A). Similar patterns were seen in beta causality: beta influences flowed to the FEF after correct outcomes and away from the FEF after incorrect outcomes (Supplemental Fig. 5B). Influences between the dlPFC and the vlPFC tended to flow from the vlPFC to the dlPFC. This directionality was consistent for all frequency bands and did not change across trial epochs (Supplemental Fig. 5). Finally, to ensure the directionality effects were not influenced by differences in signal-to-noise ratios, we recomputed Granger causality after reversing the time-series (Bastos & Schoffelen, 2016). As expected, the theta effects reversed directionality after reversing the time-series, consistent with true directionality effects (Supplemental Fig. 6).

**Figure 4.**
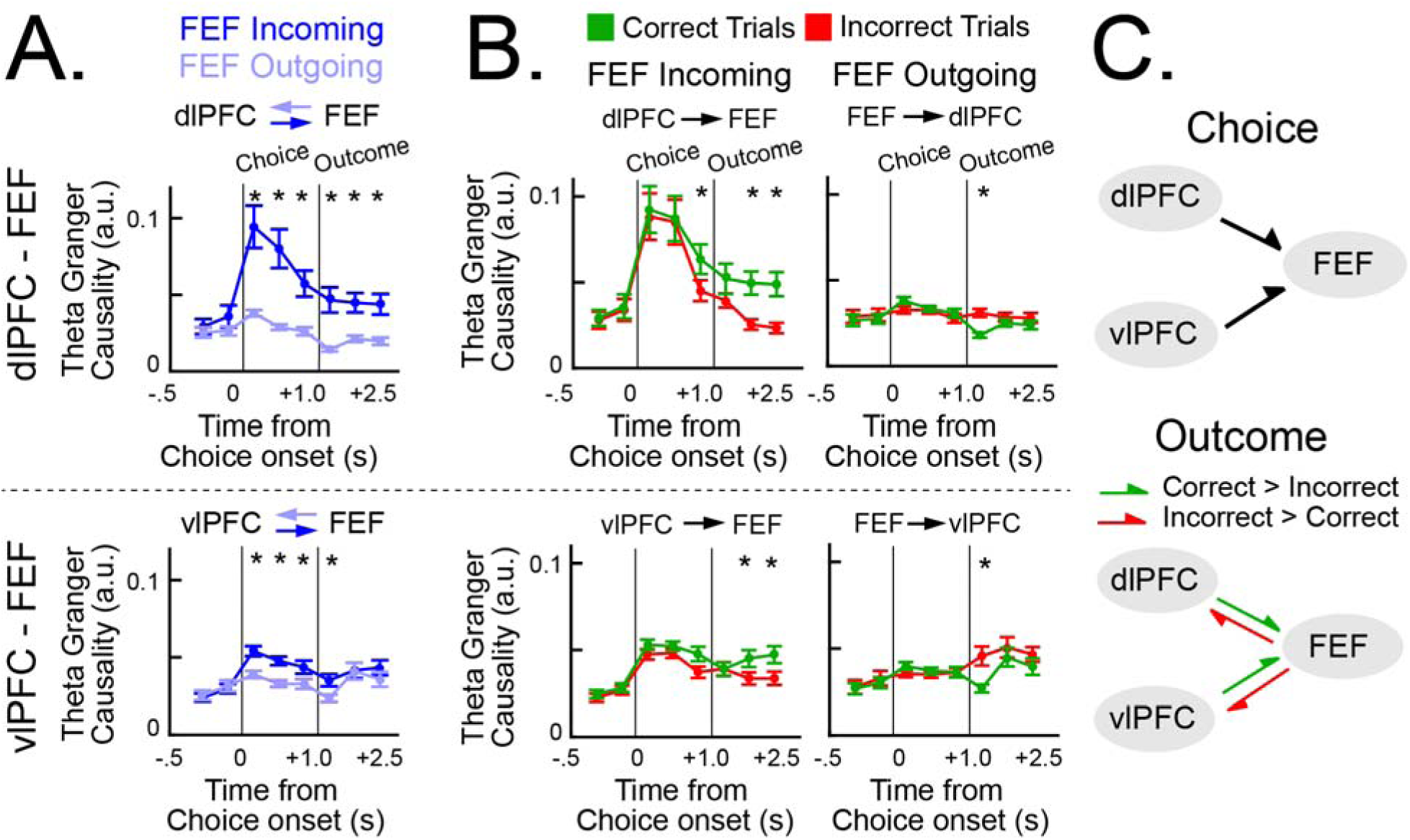
Theta Granger causality reversed directionality during the choice and outcome epochs. **A**, Average theta Granger causality between the FEF and the dlPFC/vlPFC, plotted separately by posterior-direction causality (FEF incoming; dark blue) and anterior-direction causality (FEF outgoing; light blue). * indicates timepoints in which causality was significantly different between directions (permutation test *p* < .05). **B**, Average theta Granger causality for the comparisons in **A**, separated by correct (green) and incorrect (red) trials. * indicates timepoints in which correct and incorrect causality was significantly different. **C**, Top: a summary diagram showing the effects in **A** during the choice epoch. Bottom: a summary diagram showing the effects in **B** during the outcome epoch. The color of the arrow indicates whether Granger causality was stronger after correct choices (green) or incorrect choices (red). All error bars indicate *S.E.M.* Supplemental Figure 5 shows the remaining comparisons, as well as average Granger causality in the beta and gamma frequency bands.

### Theta organized nested cross-frequency interactions

The results above indicated that theta influences were routed dynamically across the frontal areas. In the final section, we examined phase-amplitude coupling (PAC) to assess whether these theta influences affected higher-frequency oscillatory activity. For each phase-amplitude frequency pair, we computed the Modulation Index (MI), which quantifies the strength of coupling between the phase of the lower frequency and the amplitude of the higher frequency (Fig. 5A; Supplemental Fig. 7A). We examined within-area comparisons, where the same area contributed both the phase and amplitude signals. We also examined between-area comparisons, where the phase and amplitude signals were contributed by different arrays.

**Figure 5.**
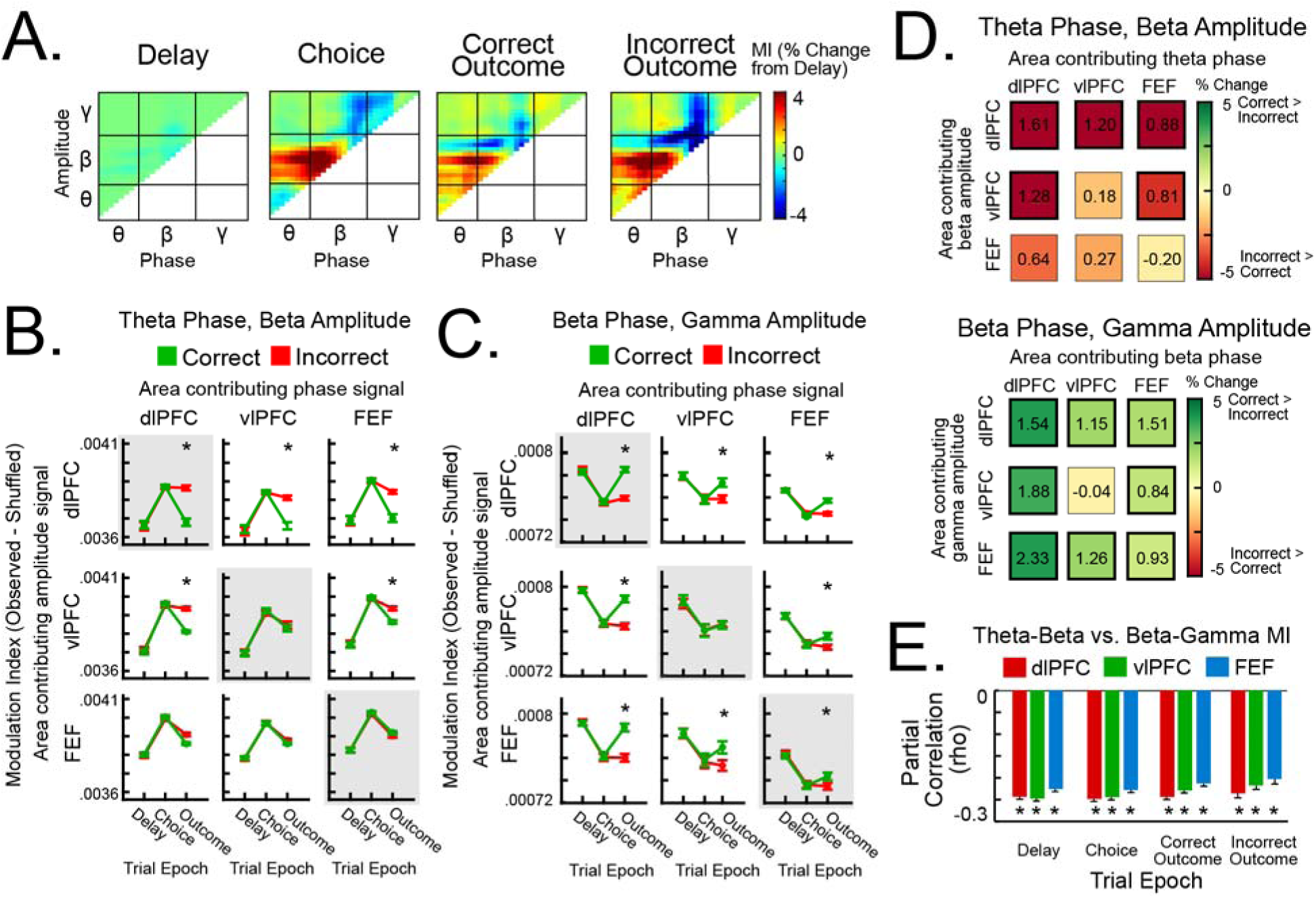
Theta-beta phase-amplitude coupling was anticorrelated with beta-gamma phase-amplitude coupling. **A,** Example “comodulogram” showing the modulation index (MI; strength of phase-amplitude coupling) for each phase-amplitude frequency pair, normalized to the Delay epoch. θ, β, γ refer to theta, beta, and gamma frequency bands, respectively. Supplemental Fig. 7A shows comodulograms for all within-area and between-area comparisons. **B-C,** Average MI for all area-area comparisons, separated by correct and incorrect trials and averaged across theta-beta frequency pairs (**B**) and beta-gamma frequency pairs (**C**). Area-area comparisons were organized according to the area that contributed the phase signal (columns) and the area that contributed the amplitude signal (rows). Within-area comparisons were on the diagonal and were outlined in gray. * indicate trials epochs in which MI was significantly different for correct and incorrect trials (permutation test *p* < .05). **D,** Summary of the results in **B-C**, during the outcome epoch. The color of each index indicates the difference in MI after correct and incorrect choices. Bolded squares indicate significant differences. The value in each square indicates Cohen’s *d* effect size. **E,** Partial correlations between single-trial theta-beta MI and beta-gamma MI. All correlations controlled for theta, beta, and gamma amplitude. * indicates significant correlation coefficients (permutation test *p* < .05). All error bars indicate *S.E.M*.

Theta phase was coupled to beta amplitude (theta-beta PAC), and beta phase was coupled to gamma amplitude (beta-gamma PAC). These were the only phase-amplitude frequency bands that reliably changed across trial epochs (Supplemental Fig. 7A). During the choice epoch, the strength of theta-beta PAC increased (Fig. 5B), and the strength of beta-gamma PAC decreased (Fig. 5C). This was consistent for within-area comparisons (Theta-beta PAC. dlPFC: Cohen’s *d* = 1.35; vlPFC: *d* = 1.24; FEF: *d* = 1.93. Beta-gamma PAC. dlPFC: *d* = 1.92; vlPFC: *d* = 1.62; FEF: *d* = 1.91), between-area comparisons (Figs. 5B&C; Theta-beta PAC: mean *d* = 1.54. Beta-gamma PAC: mean *d* = 2.02), and across NHPs (Supplemental Figs. 7B&C). During the outcome epoch, the strength of theta-beta PAC and beta-gamma PAC depended on whether the choice was correct or incorrect. Theta-beta PAC remained elevated after incorrect choices and decreased after correct choices (Fig. 5B). The difference in theta-beta PAC between correct and incorrect trials was stronger for FEF theta phase - dlPFC/vlPFC beta amplitude PAC than than opposite direction (i.e., dlPFC/vlPFC theta phase - FEF beta amplitude PAC; see Fig. 5D for Cohen’s *d* effect size). This is suggestive of an anterior directionality of theta phase during incorrect outcomes, similar to the Granger results (Fig. 4B). By contrast, beta-gamma PAC remained low after incorrect choices but increased after correct choices (Fig. 5C). This was consistent for all within-area comparisons and between-area comparisons, with the exception being vlPFC within-area PAC (see Fig. 5D for Cohen’s *d* effect size). These patterns were consistent across NHPs (Supplemental Figs. 7B&C). In sum, theta-beta PAC was stronger during choices and after incorrect outcomes, whereas beta-gamma PAC was stronger after correct outcomes.

Theta-beta PAC and beta-gamma PAC were anticorrelated on single trials. We computed partial correlations between theta-beta PAC strength and beta-gamma PAC strength during individual trials. We used partial correlations, rather than standard Pearson’s correlations, in order to control for amplitude in the theta, beta, and gamma frequency bands. For all areas, trial epochs, and NHPs, theta-beta PAC was negatively correlated with beta-gamma PAC (Fig. 5E; Supplemental Fig. 7D; mean ρ = -0.23 ± 0.004 *SEM*). This suggests that theta-beta PAC and beta-gamma PAC reflect opposing phase-amplitude configurations.

Finally, we compared the phase-amplitude relationships for theta-beta PAC (Fig. 6A) and beta-gamma PAC (Fig. 6B). For theta-beta PAC, beta amplitude tended to be highest at the falling phases of theta and lowest at the rising phases of theta (Figs. 6A&C). This phase-amplitude relationship was consistent for both within-area comparisons and between-area comparisons. By contrast, for beta-gamma PAC, gamma amplitude tended to be highest at the trough phases of beta and lowest at the peak phases of beta (Figs. 6B&D). This phase-amplitude relationship was only consistent for within-area comparisons, and not between-area comparisons. We quantified the consistency of the phase-amplitude relationship for within-area comparisons and between-area comparisons by calculating the range in amplitude across phase bins (amplitude depth; maximum minus minimum amplitude across phase bins). Figure 6E shows amplitude depth as a ratio between between-area comparisons and within-area comparisons. For all areas, this ratio was smaller for beta-gamma PAC than theta-beta PAC (Fig. 6E), suggesting that beta-gamma PAC was more local than theta-beta PAC. These patterns were consistent across trial epochs and between NHPs (Supplemental Fig. 8).

**Figure 6.**
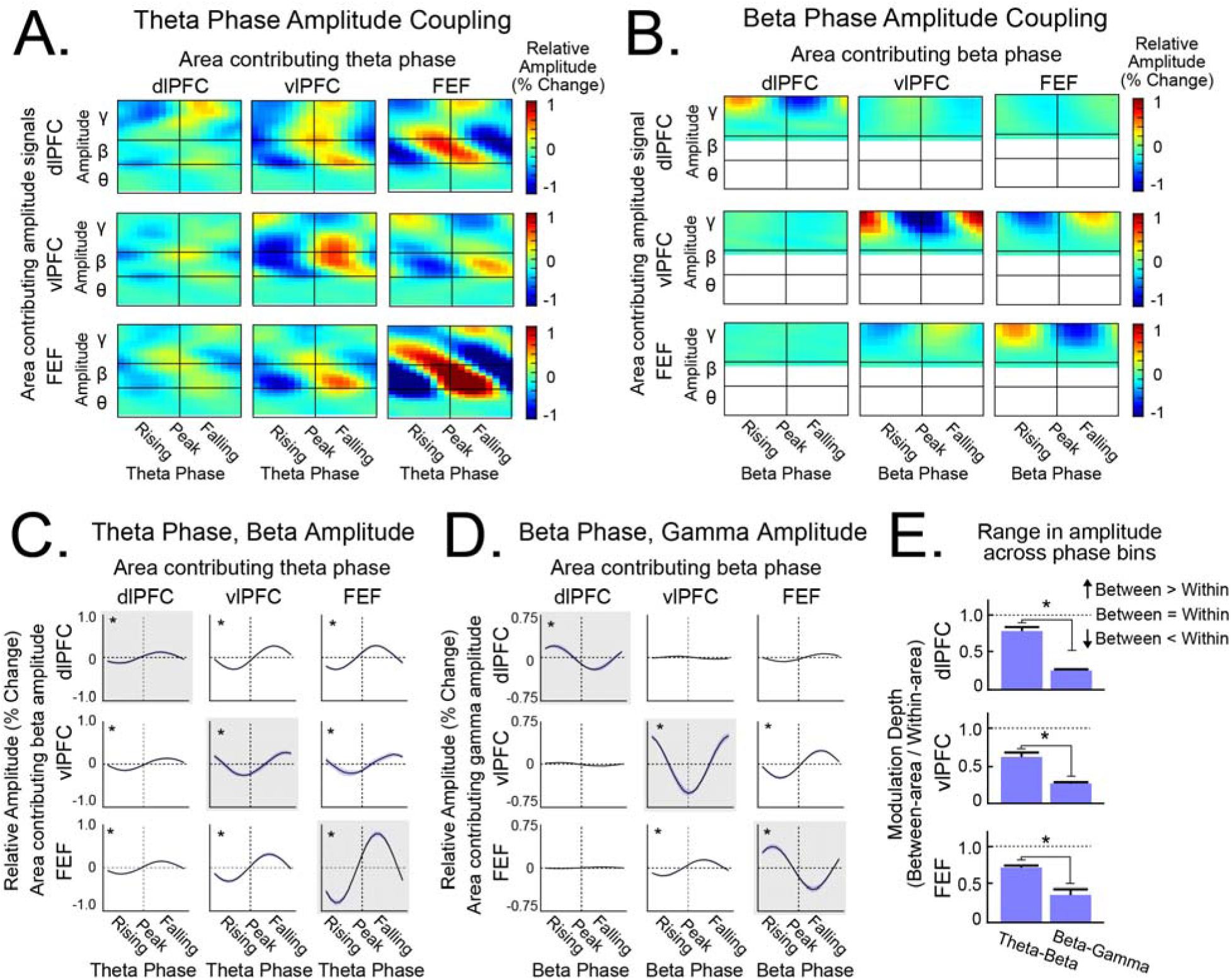
Theta-beta and beta-gamma phase-amplitude coupling peaked at different phases. **A** & **B,** Heatmaps of the relative amplitude averaged across theta phase bins (**A**) and beta phase bins (**B**). θ, β, γ refer to theta, beta, and gamma frequency bands, respectively. Area-area comparisons were organized according to the area that contributed the phase signal (columns) and the area that contributed the amplitude signal (rows). **C** & **D** Relative amplitudes averaged across phase bins for theta-beta PAC (**C**) and beta-gamma PAC (**D**). Comparisons had the same organization as (**A** & **B**). Within-area comparisons were outlined in gray. * in the left quadrant of each plot indicates that amplitude varied significantly across phase bins. See Supplemental Fig. 7 for the results in **C** & **D**, but separated by trial epoch. **E,** The range in amplitude across phase bins (“amplitude depth”), calculated as a ratio between between-area comparisons and within-area comparisons). * indicates statistical significance (permutation test *p* < .05). All error bars indicate *S.E.M*.

## DISCUSSION

Our results characterized how theta coordinated neural activity within and across frontal areas as NHPs made choices and received feedback. First, theta phase was selectively coupled to spiking information in the FEF. When the NHPs made behavioral responses, choice information in FEF spiking was coupled to LFP theta phase, in all areas. Then after receiving feedback, outcome information (and not choice information) became coupled to theta phase, in all areas. Second, theta influences were routed in opposite directions depending on feedback. During behavioral choices and following correct outcomes, theta influences flowed in the posterior direction from the dlPFC/vlPFC to the FEF. Following incorrect outcomes, theta influences reversed and flowed in the anterior direction from the FEF to the dlPFC/vlPFC. Third, theta organized nested cross-frequency, phase-amplitude interactions. Theta phase was coupled to beta amplitude (theta-beta PAC), and beta phase was coupled to gamma amplitude (beta-gamma PAC). Theta-beta PAC was strong during choices and after incorrect outcomes, and beta-gamma PAC was strong after correct outcomes. Together, our results provide a multi-level account of how theta dynamically coordinates activity in the frontal cortex.

Theta seemed to function as a gating mechanism to select relevant spiking information. Many reports have shown that theta phase couples to spiking activity (O’Keefe & Recce, 1993; Rutishauser et al., 2010; Reddy et al., 2021; Benchenane et al., 2010; Han et al., 2026). Fewer studies have shown that spiking information, as opposed to spike rates, also couples to LFP phase (Voloh et al., 2020; Kayser et al., 2009). Our results extend this literature by showing that the type of information coupled to theta phase depended on what was immediately relevant. During the choice and outcome epochs, spiking activity carried information about multiple task variables simultaneously, as they often do (Rigotti et al., 2013; Mante et al., 2013; Cromer et al., 2010). However, only currently relevant information was coupled to theta phase: choice information during choices, and outcome information following feedback. Theta phase thus seemed to organize which information was available. This provides a mechanism by which information can be dynamically reconfigured during changing task demands.

This gating mechanism may depend on an interaction of excitation and inhibition. The optimal phase of decoding spiking information was offset from the peak phase of spike-field coupling. This offset is consistent with previous reports (Kayser et al., 2009; Voloh et al., 2020; Schaefer et al., 2006) and helps rule out the possibility that information decoding was higher simply because spiking activity was high. Furthermore, the timing hints at a more complex interaction between excitation and inhibition. At the peak of theta, overall spike rates increased. Then during the falling phases of theta, activity waned, but information increased. This is suggestive of inhibitory mechanisms (e.g., interneurons; Rao et al., 2000; Lewis et al., 2012) that “shape” information from overall activation. Indeed, beta amplitude, which has been implicated in inhibitory control (Lundqvist et al., 2023; Chen et al., 2026), also peaked during the falling phases of theta.

The FEF emerged as a dynamic hub. During behavioral choices, theta influences flowed into the FEF from the dlPFC and the vlPFC. This replicates previous studies (Babapoor-Farrokhran et al., 2017) and is consistent with a role of the FEF in relaying top-down information from the prefrontal cortex to the visual cortex in order to direct spatial attention (Moore & Armstrong, 2003; Moore & Fallah, 2001; Schafer & Moore, 2007). These results are also consistent with a more recent study showing that FEF theta is critical for the readout of working memory content (Han et al., 2026). However, after incorrect choices, theta influences reversed directionality and flowed anteriorly back to the dlPFC/vlPFC. This reversal suggests that the FEF functioned as a critical node in the evaluation of choices and outcomes (Teichert, Yu, & Ferrera, 2014; Shteyn & Olson, 2024; Ding & Hikosaka, 2006) and was not a simple relay of top-down information. Furthermore, only FEF spiking information was coupled to theta phase. Spiking activity in the dlPFC/vlPFC carried choice and outcome information, but these were not coupled to LFP phase. Together, these results suggest that the FEF played a privileged role in selecting and broadcasting choices and errors.

Our observations of theta-beta PAC and beta-gamma PAC were unexpected, as many studies have focused only on coupling between theta phase and gamma amplitude (Tort et al., 2010; Lisman & Jensen, 2013; Belluscio et al., 2012; Saint Amour di Chanaz et al., 2023; Canolty et al., 2006; Tamura et al., 2017; Spyropoulos, Bosman, & Fries, 2018; Bonnefond, Kastner, & Jensen, 2017; Voloh et al., 2015; Lisman & Idiart, 1995). In our study, theta-gamma PAC was present, but did not reliably change across trial epochs or by trial outcome. This suggests that theta had a more indirect influence on gamma, via beta (Bastos et al., 2015; Lundqvist et al., 2016; Lundqvist et al., 2018). In fact, theta-beta and beta-gamma PAC were anticorrelated and seemed to reflect opposing network configurations. For theta-beta PAC, the phase-amplitude relationship was consistent for within-area and between-area comparisons, suggesting that theta-beta PAC may be more important for more long-distance routing. By contrast, for beta-gamma PAC, the phase-amplitude relationship was only consistent for within-area comparisons, suggesting that beta-gamma PAC may be more important for local coupling. Beta, as the intermediate frequency, may mediate between the long-distance routing of theta-beta PAC and the local maintenance of beta-gamma PAC. This would be consistent with beta’s role in top-down control (Buschman & Miller, 2007; Bastos et al., 2015; Lundqvist et al., 2024; Spitzer & Haegens, 2017).

Taken together, our results suggest that theta functions as a “master clock” that coordinates network dynamics in the frontal cortex. This coordination occurred at multiple levels. Theta gated spiking information, routed influences between subareas, and organized higher-frequency oscillations. For each of these mechanisms, theta’s role was not static across the trial, but shifted dramatically from choices to outcomes. Together, these findings emphasize theta’s role in governing flexible cognition.

## MATERIAL & METHODS

### Task procedures

All task procedures have been previously described (Loonis et al., 2017). Briefly, non-human primates (NHPs; two adult rhesus macaques; NHP 1: female, 6 years, 6.5 kg; NHP 2: male, 9 years, 13 kg) learned to categorize dot patterns. Novel category prototypes were generated before each session. These prototypes were designed such that task difficulty was roughly equivalent across sessions (see Loonis et al., 2017 for details). Category exemplars were created by distorting the dot positions of the prototype (Posner & Keele, 1968; Distortion level 1). The number of category exemplars increased progressively as the NHPs learned the categories. All procedures followed the guidelines according to the Massachusetts Institute of Technology Committee on Animal Care and the National Institutes of Health.

The NHPs started each trial by fixating on a central fixation point (2.5 visual degrees) for 0.5 seconds (Fig. 1B; fixation epoch). Then, a category exemplar (7 visual degrees x 7 visual degrees) was presented at the center of the screen for 1.0 second (sample epoch). After a variable working memory delay (delay epoch; 0.85 seconds to 1.25 seconds), two new exemplars were presented on the left and right sides of the screen (choice; 9 visual degrees from the center). One choice exemplar came from the same category as the sample exemplar (i.e., its “match” category). The other exemplar came from the other, “non-match” category. The NHPs were trained to fixate on the match exemplar for at least 0.7 seconds. Visual feedback was given after each choice. After correct choices, the chosen exemplar turned from white to green, and the NHPs received a juice reward (simultaneously). After incorrect choices, the chosen exemplar turned red, juice was withheld, and the NHPs were given a “timeout” (5–16 seconds) before the start of the next trial.

The NHPs completed multiple blocks of trials during each session. The number of unique category exemplars increased exponentially across blocks (i.e., 2^block^ number of exemplars). The NHPs progressed to the next block after reaching a learning criterion (i.e., >80% accuracy on the previous twenty trials and >70% accuracy on the previous ten trials for each of the four main conditions: Category ‘A’ on the left, Category ‘A’ on the right, Category ‘B’ on the left, Category ‘B’ on the right). For the current analysis, we excluded sessions in which the NHPs did not reach at least training block six (i.e., 10 sessions were excluded out of 76 total sessions). We only analyzed trials during and after training block three, to focus on post-learning neural coding.

### Electrophysiological Recordings

All experimental sessions were controlled by custom MATLAB software using PsychToolbox. Stimuli were presented on a LCD screen (ViewSonic VG2401mh 24_′′_ Gaming Monitor). Eye movements and pupil size were monitored using EyeLink II sampled at 1000 Hz. Electrode arrays (Blackrock Cereport; 8 x 8 grid with 400 µm spacing; 1 mm length) were implanted into the dorsolateral prefrontal cortex (dlPFC), ventrolateral prefrontal cortex (vlPFC), and the frontal eye fields (FEF). Figure 1C shows the approximate position of each array in respect to the principal and arcuate sulci.

Signals were sampled at 30 kHz, band-passed between 0.3 Hz and 7.5 kHz using a Butterworth filter, and digitized at a resolution of 16 bits, 250 nV/bit. LFPs were recorded with a low-pass 250 Hz Butterworth filter, sampled at 1 kHz, referenced to ground, and AC-coupled. Evoked potentials averaged from correct and incorrect trials were subtracted from each individual electrode. Multiunit activity was recorded for each channel. Channels with a grand average firing rate less than 0.5 Hz were excluded from all analyses.

Time-resolved phase and amplitude information were calculated for thirty frequencies, ranging from 4 Hz to 90 Hz along a log scale. For each frequency, the raw LFP time series was bandpass filtered (Butterworth filter, 4^th^ order; filter centered on that frequency). Then, the Hilbert transform was computed on the bandpass-filtered data. Phase and amplitude were determined as the arctan and absolute value of the complex signal, respectively. Frequency bands were defined using these frequency ranges: theta (4–10 Hz), beta (15–30 Hz) and gamma (40–90 Hz).

### Phase-Amplitude Coupling

Phase-amplitude coupling (PAC) was computed to examine cross-frequency interactions. Coupling was computed between phase-amplitude frequency pairs in which the “amplitude” frequency was higher than the “phase” frequency (i.e., the upper triangle in the full phase-amplitude matrix; see Fig. 5A for an example). Trials were divided into 500 ms non-overlapping windows. This window size was selected to ensure all phase-amplitude frequency pairs included data from at least two complete phase cycles. For each window, the phase of the lower frequency was binned into eighteen equal-sized bins (i.e., twenty degrees per bin). The amplitude of the higher frequency was averaged for each phase bin, separately. The binned amplitudes were normalized by dividing each amplitude by their sum across phase bins. This resulted in a set of normalized amplitudes for each trial, trial window, and phase-amplitude frequency pair.

The strength of phase-amplitude coupling was determined by calculating the Modulation Index (MI; Tort et al., 2009). Intuitively, MI quantifies the extent to which amplitude is non-uniformly distributed across phase bins. A larger MI value would indicate that the amplitude in the higher frequency varied according to the phase of the lower frequency. MI was computed as follows:

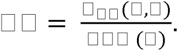

□_□□_ was the Kullback-Leibler (KL) divergence between vectors *P* and *U*, where *P* contained the observed normalized amplitudes across the *N* phase bins, and *U* contained the theoretical amplitudes when assuming a uniform distribution (i.e., 1/N). Specifically, □_□□_ was calculated as:

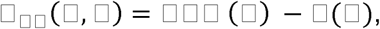

where *H* was the Shannon entropy across phase bins, defined as:

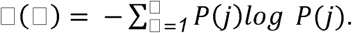

MI values range from 0 to 1, where 0 would indicate that the amplitudes were perfectly uniform across phase bins, and 1 would indicate that the amplitudes were perfectly clustered at a single phase bin. MI values were computed for each trial, trial window, and phase-amplitude frequency pair separately and then averaged across trials.

To ensure the MI estimates were not influenced by spurious correlations between the phase and amplitude signals, we computed a null distribution of shuffled MI values. Specifically, we added a circular shift (with a random offset) to the LFP phase signal before binning the amplitudes and calculating MI. This was repeated one hundred times. We subtracted off the average value of these shuffled iterations from the observed MI value.

We examined both within-area comparisons, where the phase signal and amplitude signals were computed from the same array, and between-area comparisons, where the phase and amplitude signals were computed from separate arrays. The phase signal was determined by computing the circular mean of phases across all electrodes within that array. The PAC results were qualitatively similar when, instead of averaging phase across electrodes, phase was determined by a “representative” electrode (i.e., the center-most electrode in the array).

### Granger Causality

Directional influences between LFPs across areas were measured using spectral Granger causality (sGC; Geweke, 1982). Like traditional Granger causality, sGC quantifies how variance in one time-series can be explained by the past values of another time-series, beyond the variance that can be explained using that time-series’s own past values. Unlike traditional Granger causality, sGC expresses these influences in the spectral (frequency) domain and does not rely on fitting a specific autoregressive model. We computed sGC between each pair of between-area electrodes in 500 ms non-overlapping time windows. Directionality was assessed by comparing sGC in each direction (i.e., forward-direction vs. backward-direction). To directly compare sGC between conditions with unequal numbers of trials (i.e., correct vs. incorrect), we subsampled trials according to the condition with a lower trial count. To ensure that results reflected true directional effects, rather than between-area differences in signal-to-noise ratio, we confirmed that estimated directionalities reversed when signals were time-reversed (Supplementary Fig. 6). All analyses were conducted using the *FieldTrip* toolbox in MATLAB.

### Partial Correlations

Partial correlations were computed to examine whether the strength of phase-amplitude coupling was correlated during single trials. We used partial correlations, rather than standard Pearson’s correlations, to control for potential sources of shared variance between variables (i.e., theta, beta, and gamma amplitude). For each trial epoch and area, we computed single-trial estimates for theta-beta MI, beta-gamma MI, theta amplitude, beta amplitude, and gamma amplitude. Each variable was normalized by computing z-scores across all trials. Partial correlations were computed between theta-beta MI and beta-gamma MI, after controlling for theta, beta, and gamma amplitude. Correlation coefficients were averaged across sessions using Fisher’s z-transformation. All correlations were computed using the *partialcorr* function in MATLAB.

### Spike Decoding

Support Vector Machine (SVM) classifiers (Cortes & Vapnik, 1995) were trained with spiking activity from each area to decode three task variables: 1) the category membership of the sample exemplar (category information; category ‘A’ or category ‘B’), 2) the location of the category “match” exemplar during the choice epoch (i.e., choice information; left or right), and 3) the feedback given to the NHPs (outcome information; correct or incorrect). Spiking activity was binned into 50 ms non-overlapping windows across the trial epochs. Spiking activity in each electrode was normalized by computing z-scores across all trials and timepoints. Each classifier was trained using 80% of trials, and then cross-validated on the held-out 20% of trials. We ensured the training set had an equal number of trials in each condition (e.g., correct vs. incorrect). Classifier accuracies for the held-out trials were averaged across one hundred iterations, where the trials used in the training set were randomized during each iteration. All model fits were computed using the *fitcsvm* function in MATLAB.

We also tested whether spiking information varied across LFP phases. We divided each trial into 500 ms non-overlapping windows. Within each time window, we divided the LFP phase into 45-degree bins (i.e., eight total bins). We summed (and normalized) spiking activity using timepoints from each phase bin, separately. We trained SVM classifiers to decode task variables using the procedures described above. This resulted in a curve of classifier accuracies across phase bins for each LFP frequency. The classifier accuracies were normalized by dividing by their sum across phase bins. Phase-locking values (PLVs) were computed to quantify the extent to which classifier accuracies changed across phase bins. Specifically, PLVs were defined as:

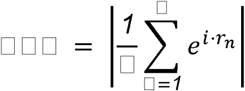

where *N* is the number of phase bins, and □_□_ is the classifier accuracy for phase bin *n*. These PLVs were compared against a null distribution of PLVs, in which the classifier accuracies were shuffled across phase bins before computing PLV. This was repeated 1,000 times, and the average PLV from these shuffled iterations was subtracted off from the observed PLV.

We examined both within-area comparisons, where spiking activity and LFP phases were computed from the same array, and between-area comparisons, where spiking activity and LFP phase were computed from separate arrays. LFP phase was computed according to the circular mean of phases from all electrodes in the array. Results were qualitatively similar if, instead of averaging across electrodes, phase was determined from a “representative” electrode (i.e., the center-most electrode in the array).

### Spike-Field Coupling

We also examined coupling between spike rates and LFP phase. We divided trials into 500 ms non-overlapping windows. For each trial, LFP phase was divided into 18 bins (20 degrees per phase bin). Then for each neuron, we computed the number of spikes that fell within each phase bin. We examined within-area comparisons, where the spiking activity and LFP phase were computed from the same array. We also examined between-area comparisons, where the spiking activity and LFP phase were computed from different arrays. LFP phase was computed as the circular mean from all electrodes in that array. Results were similar if, rather than averaging across electrodes, LFP phase was determined from a “representative” electrode (i.e., the center-most electrode).

### Statistical Analysis

For continuous neural measures that we could not assume were normally distributed, we used nonparametric statistics. All measures were averaged across electrodes within a given array, and one estimate was used per session (i.e., *n* = 66 total sessions). Randomizations were performed across all samples and were resampled 10,000 times to obtain a *p* value. Cohen’s *d* was computed to quantify the effect size of the main effects. Measures were averaged across timepoints pertaining to each trial epoch (e.g., theta power during the choice epoch for correct and incorrect trials). The effect size of this difference (e.g., correct – incorrect) was calculated as the mean difference across sessions, divided by the standard deviation across sessions.

## Acknowledgements

Army Research Office W911NF2410228; Freedom Together Foundation; The Picower Institute for Learning and Memory; Office of Naval Research MURI N00014-23-1-2768

## Author Contributions

*Conceptualization:* MBB, SLB, RFL, EKM. *Methodology:* MBB, SLB, RFL, EKM. *Formal analysis:* MBB. *Investigation:* RFL. *Data Curation:* RFL, SLB. *Writing-Original Draft:* MBB, SLB, EKM. *Writing- Review & Editing:* MBB, RFL, SLB, EKM. *Visualization:* MBB. *Supervision:* EKM. *Project administration:* EKM, MBB. *Funding acquisition:* EKM

## SUPPLEMENTAL FIGURES

**Supplemental Figure 1.**
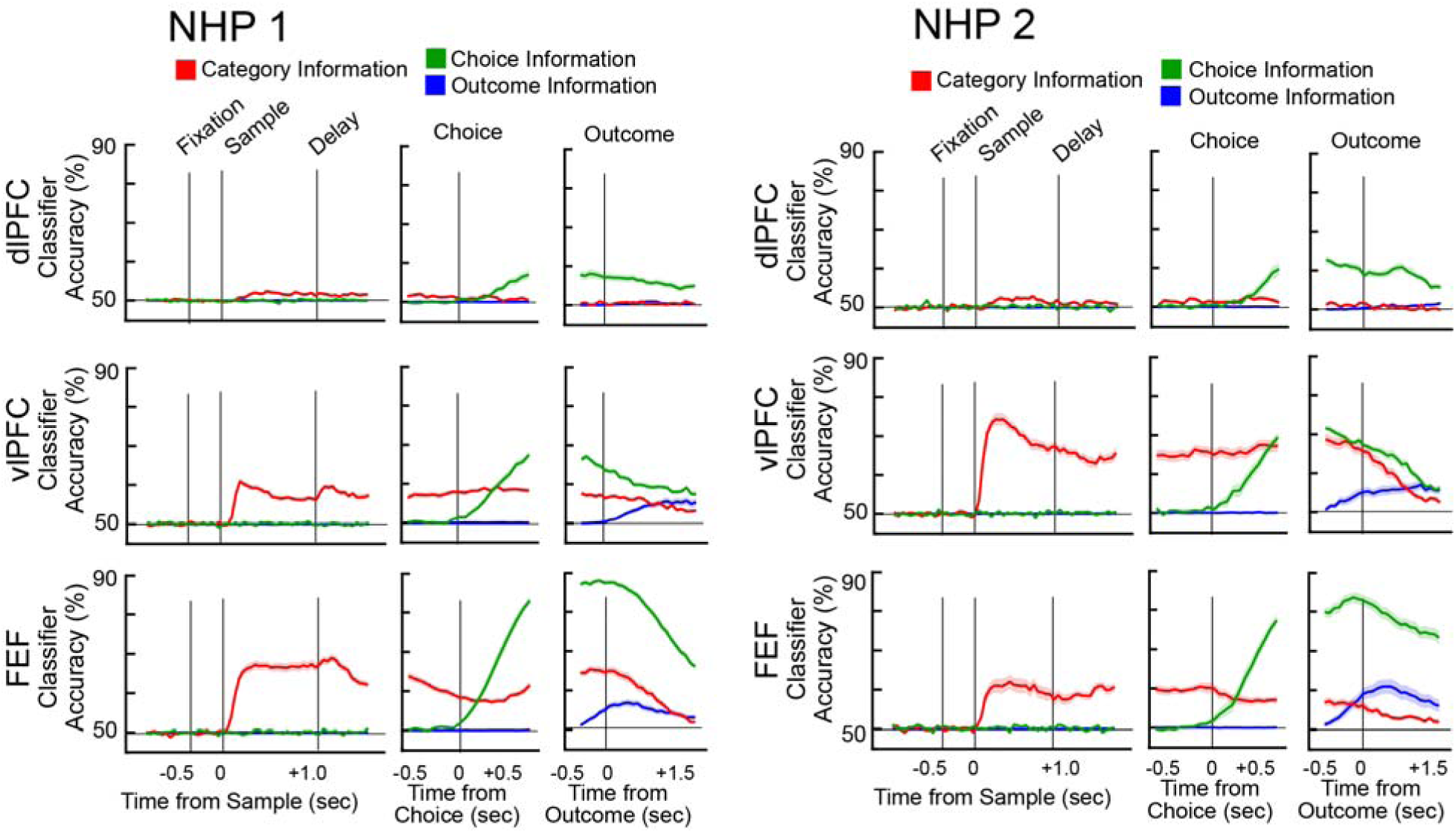
Spiking activity carried information about task variables (results related to Fig. 2A). Average SVM classifier accuracies in each area and task variable, separated for NHP 1 (left) and NHP 2 (right). The color of each plot indicates the type of information being decoded. All error bars indicate *S.E.M.* Choice and outcome epoch results are replotted from Fig. 2A; this figure additionally plots information during the sample and delay epochs.

**Supplemental Figure 2.**
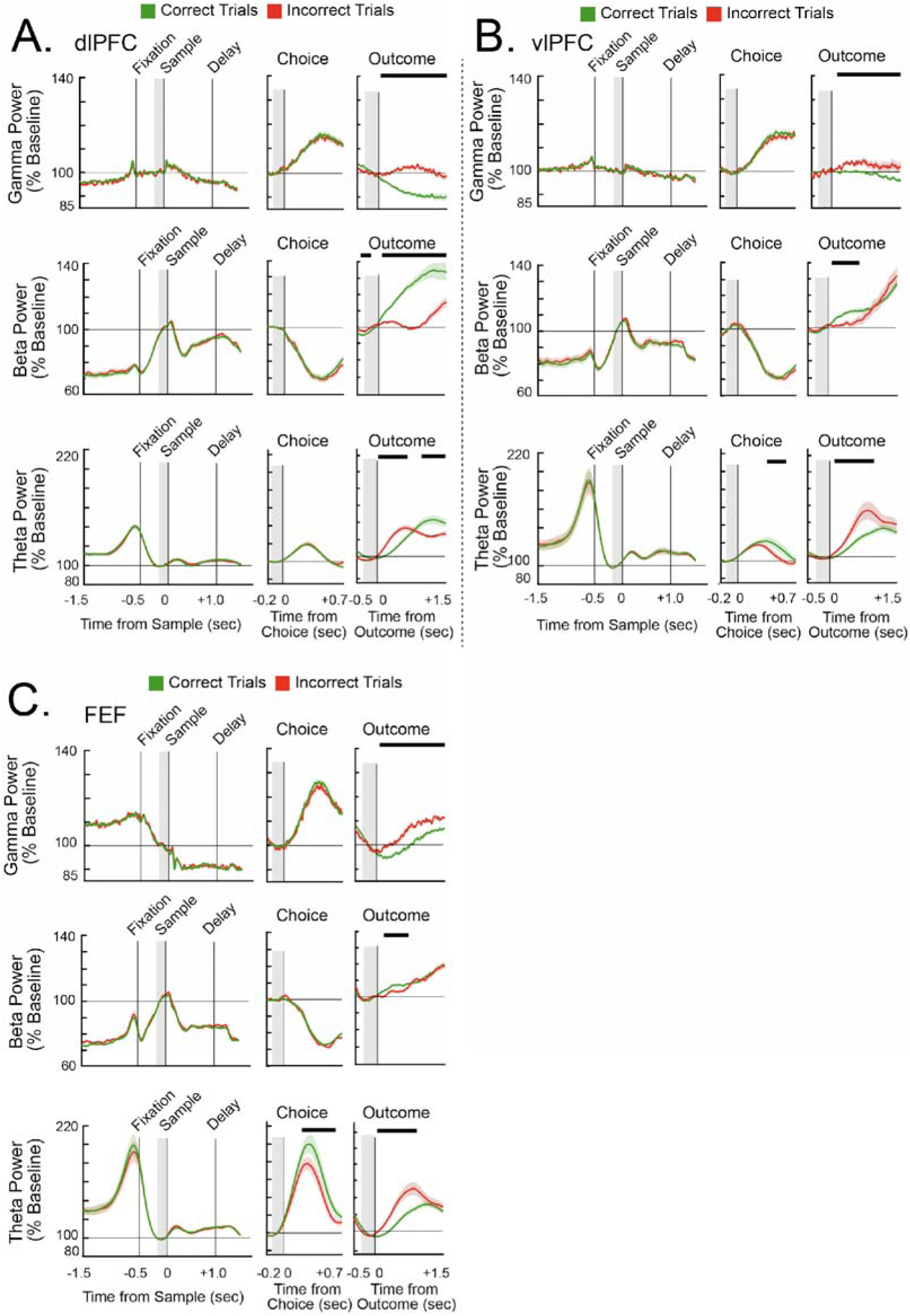
Average power reflected choices and outcomes (results related to Figs. 2B&C). Power in the dlPFC (**A**), vlPFC (**B**), and FEF (**C**), averaged in gamma (top), beta (middle), and theta (bottom) frequency bands, and separated by correct (green) and incorrect (red) trials. Bars above each plot indicate timepoints in which power was significantly different between correct and incorrect trials. All error bars indicate *S.E.M*.

**Supplemental Figure 3.**
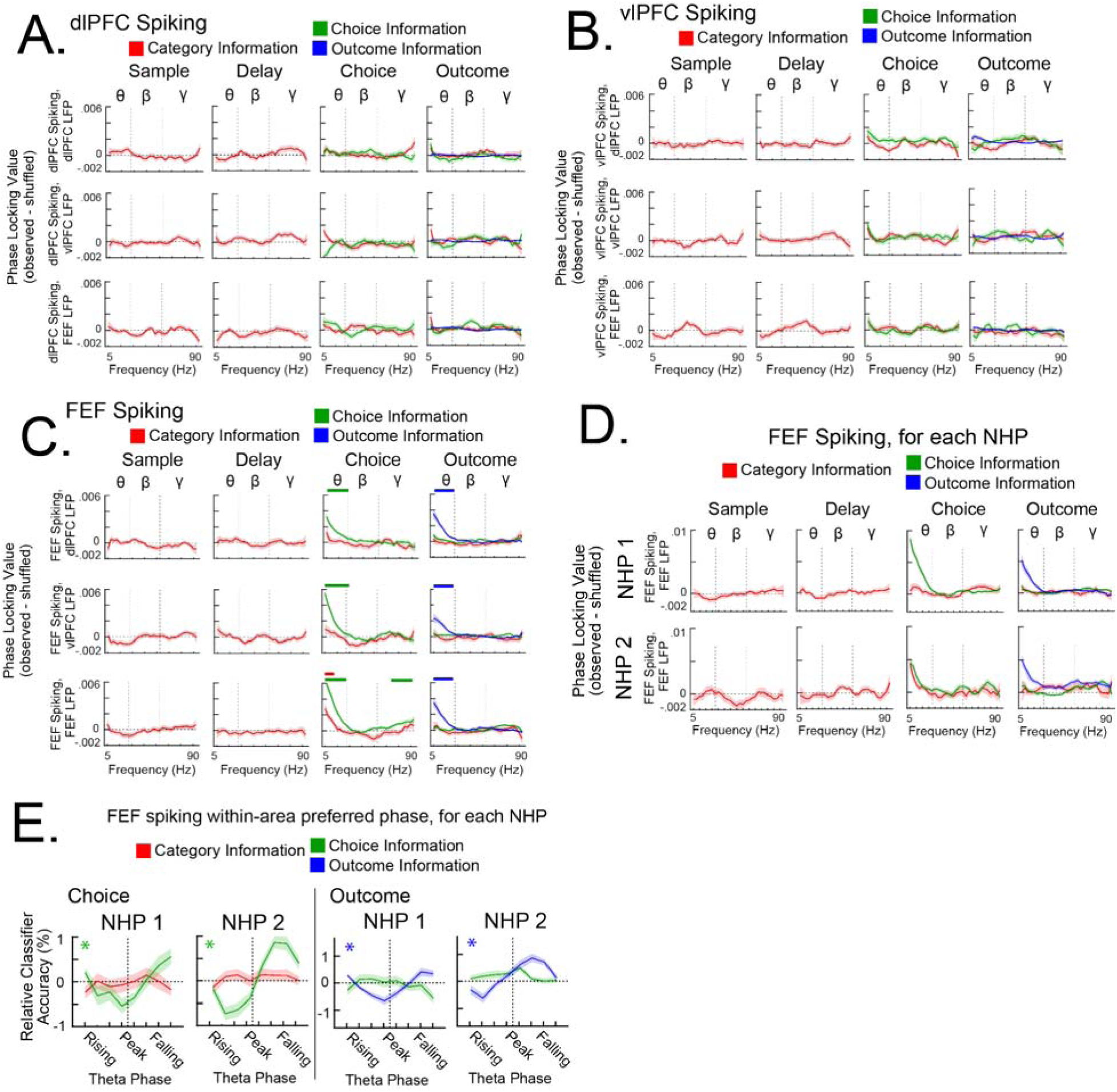
Coupling between binned LFP phase and spiking information in each area (related to Figs. 3A-D). **A-C**, Phase-locking values quantified whether classifier accuracies varied with LFP phase. All within-area and between-area comparisons for dlPFC spiking (**A**), vlPFC spiking (**B**), and FEF spiking (**C**). Colored bars above each plot indicate frequencies in which spiking information was significantly coupled to LFP phase. **D**, Within-area FEF results (bottom row of **C**), separated by NHP 1 (top) and NHP 2 (bottom). **E**, Average classifier accuracies across theta phase bins, separated by NHP 1 (left) and NHP 2 (right). All error bars indicate *S.E.M.* θ, β, γ refers to theta, beta, and gamma frequency bands, respectively.

**Supplemental Figure 4.**
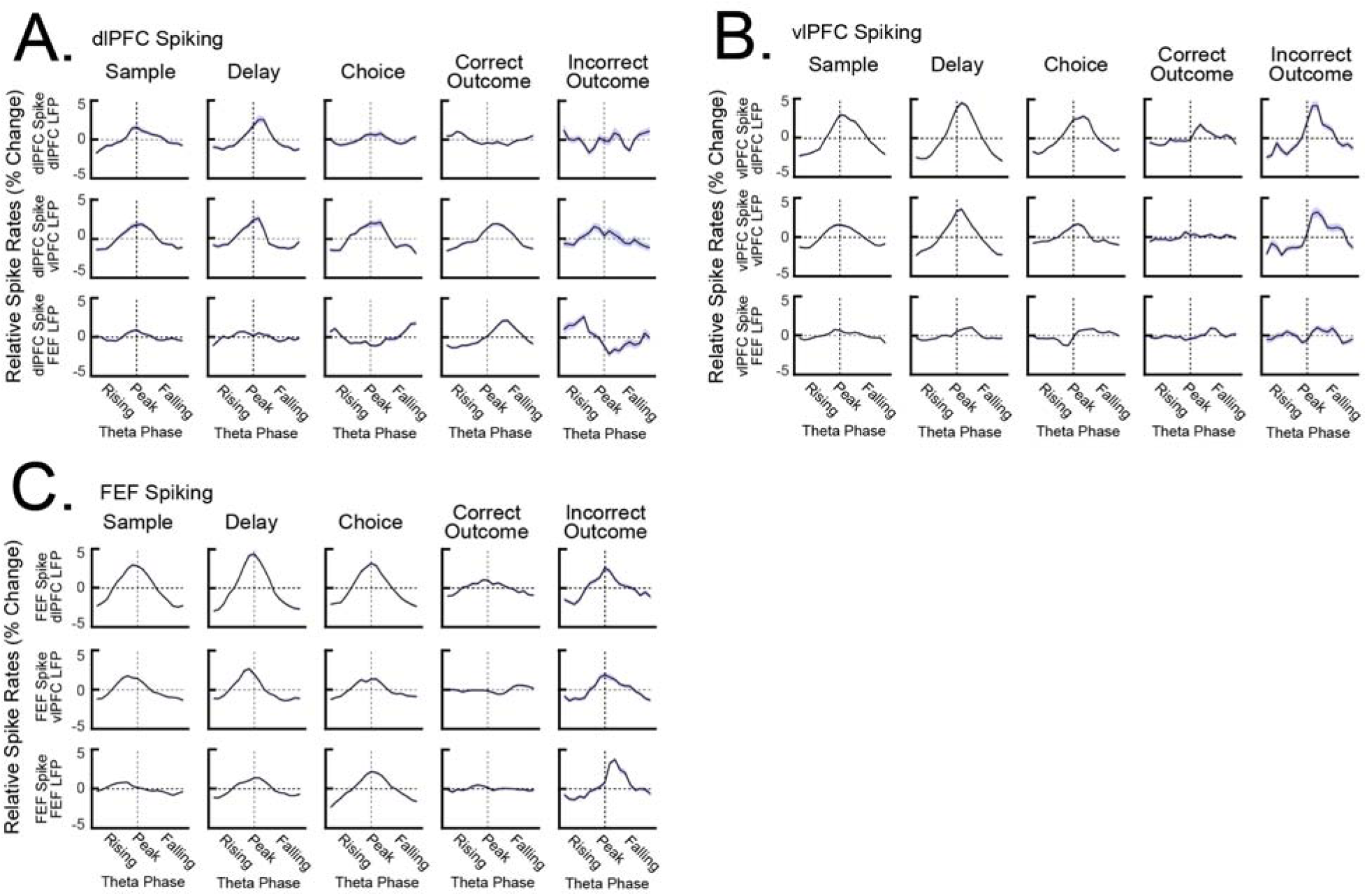
Spike-field coupling (results related to Figs. 3E&F). Relative spike rates were averaged across theta phase bins. All within-area comparisons and between-area comparisons for dlPFC spiking (**A**), vlPFC spiking (**B**), and FEF spiking (**C**). Results were separated by trial epoch. All error bars indicate *S.E.M*.

**Supplemental Figure 5.**
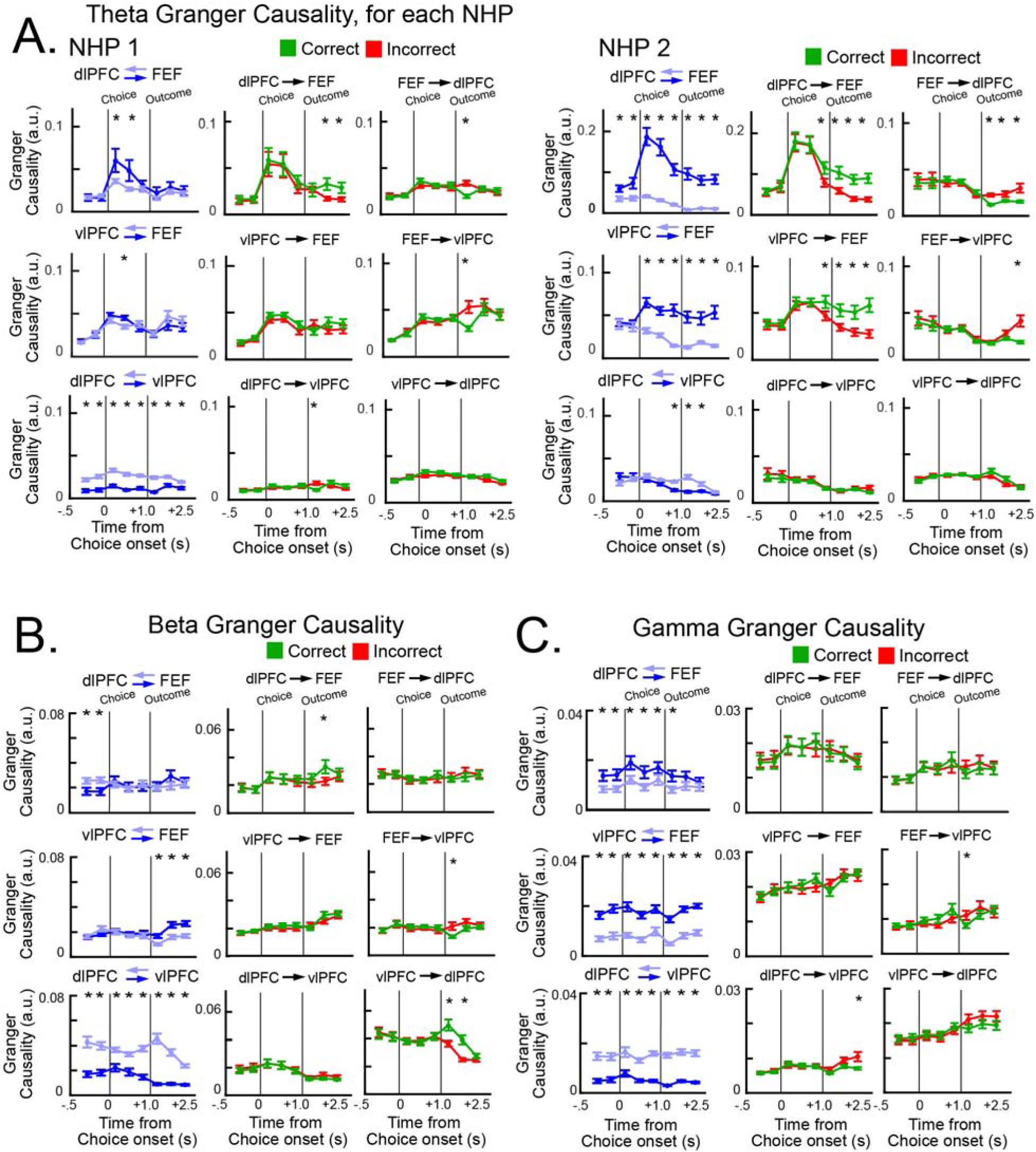
Average Granger causality for each frequency band and NHP (related to Fig. 4). **A**, Average theta causality for NHP 1 (left) and NHP 2 (right). **B-C**, Average Granger causality in the beta (**B**) and gamma (**C**) frequency bands. * indicates timepoints in which causality was significantly different between comparisons. All error bars indicate *S.E.M*.

**Supplemental Figure 6.**
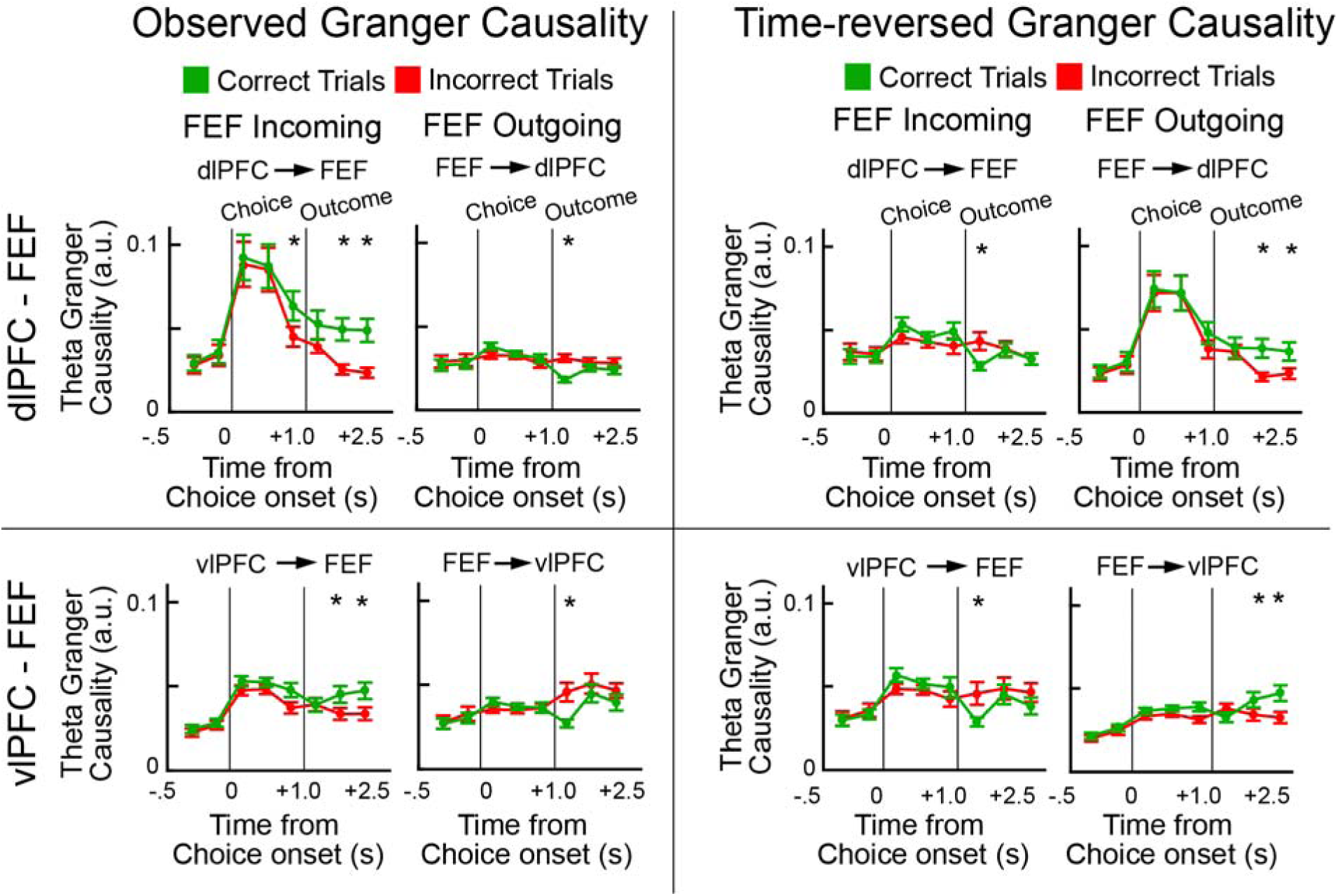
Time-reversed theta Granger causality (results related to Fig. 4). We recomputed Granger causality after reversing the time-series as a control analysis to test whether the directionality effects were influenced by possible differences in the signal-to-noise ratio across arrays. Left: the observed theta Granger causality, separated by correct and incorrect trials. Right: time-reversed theta Granger causality, separated by correct and incorrect trials. Importantly, the directionality of effects reversed when the time-series were reversed. This is consistent with “true” directionality effects. * indicates statistical significance (permutation test *p* < .05). All error bars indicate *S.E.M*.

**Supplemental Figure 7.**
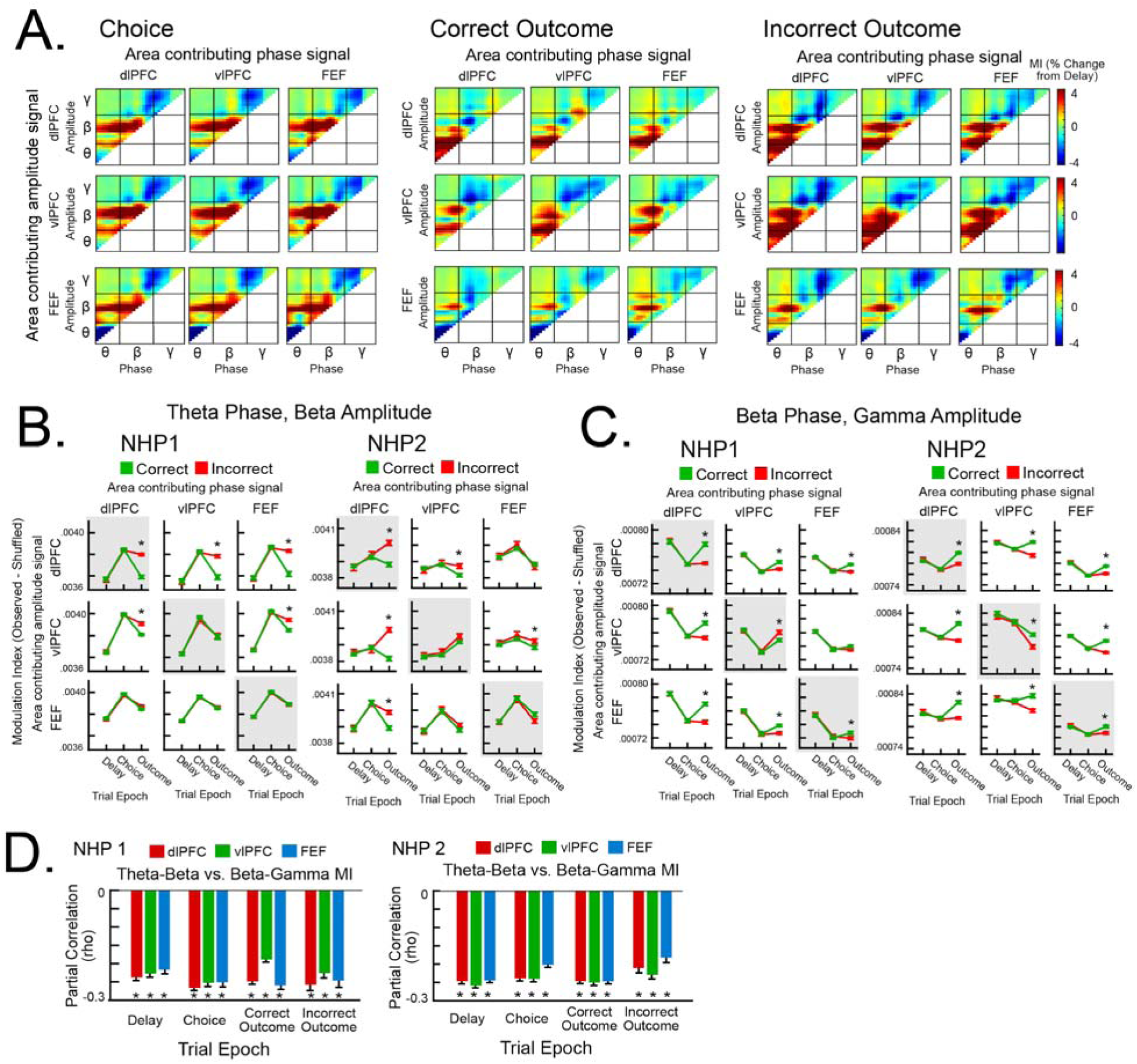
Phase-amplitude coupling (PAC; related to Figure 5). **A, “**Comodulograms” showing the Modulation Index (MI) between each phase-amplitude frequency pair, normalized to the delay epoch. Area-area comparisons were organized according to the area that contributed the phase signal (columns) and the area that contributed the amplitude signal (rows). θ, β, γ refers to theta, beta, and gamma frequency bands, respectively. **B-C**, Average MI for theta-beta PAC (**B**) and beta-gamma PAC (**C**), separated for NHP 1 (left) and NHP 2 (right). Area-area comparisons were organized in the same way as **A**, with within-area comparisons outlined in gray. * indicates timepoints in which MI was significantly different between correct and incorrect trials. **D**, Partial correlations between single trial theta-beta MI and beta-gamma MI, separated for NHP1 (left) and NHP 2 (right). * indicates correlation coefficients that were significantly different from 0. All error bars indicate *S.E.M*.

**Supplemental Figure 8.**
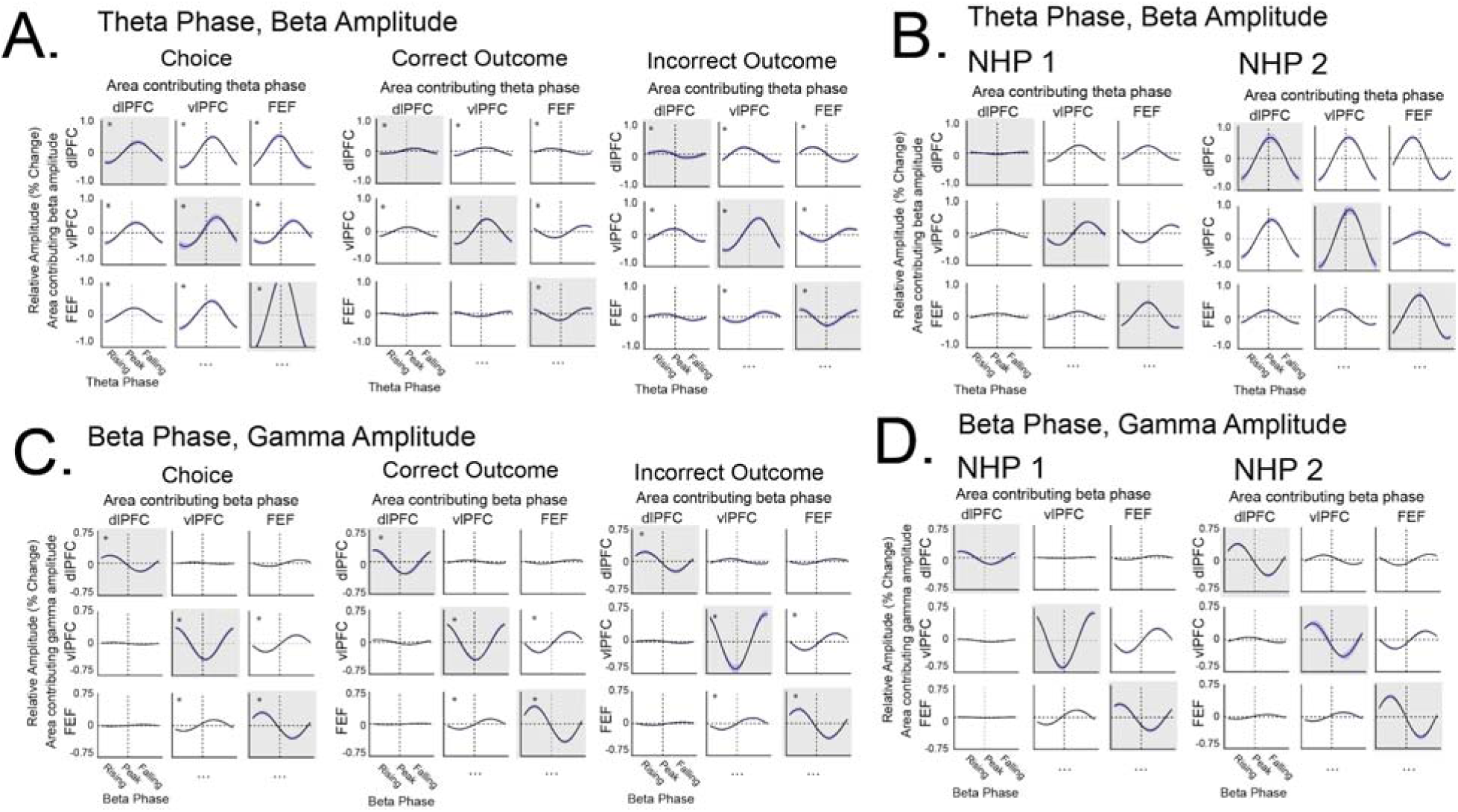
Average phase-amplitude relationships (PAC; related to Figure 6). Area-area comparisons were organized by the area that contributed the phase signal (columns) and the area that contributed the amplitude signal (rows), with within-area comparisons outlined in gray. **A**, Average theta-beta PAC for each trial epoch. **B**, Average theta-beta PAC, separated by NHP. **C**, Average beta-gamma PAC for each trial epoch. **D**, Average beta-gamma PAC, separated by NHP. All error bars indicate *S.E.M*.

## Notes

**Conflict of interest:** The authors declare no competing financial interests.

### Competing Interest Statement

The authors have declared no competing interest.

